# Locus Suicide Recombination actively occurs on the functionally rearranged IgH allele in B-cells from inflamed human lymphoid tissues

**DOI:** 10.1101/430215

**Authors:** Iman Dalloul, François Boyer, Zeinab Dalloul, Amandine Pignarre, Gersende Lacombe, Thierry Fest, Fabrice Chatonnet, Céline Delaloy, Anne Durandy, Robin Jeannet, Emilie Lereclus, Hend Boutouil, Jean-Claude Aldigier, Sophie Peron, Jeanne Cook-Moreau, Michel Cogné

## Abstract

B-cell activation yields abundant cell death in parallel to clonal amplification and remodeling of immunoglobulin (Ig) genes by activation-induced deaminase (AID). AID promotes affinity maturation of Ig variable regions and class switch recombination (CSR) in mature B lymphocytes. In the IgH locus, these processes are under control by the 3’ regulatory region (3’RR) super-enhancer, a region demonstrated in the mouse to be both transcribed and itself targeted by AID-mediated recombination. Alternatively to CSR, IgH deletions joining Sμ to “like-switch” DNA repeats that flank the 3’ super-enhancer can thus accomplish so-called “locus suicide recombination” (LSR) in mouse B-cells. We now show that AID-mediated LSR also actively occurs in humans, and provides an activation-induced cell death pathway in multiple conditions of B-cell activation. LSR deletions either focus on the functional IgH allele or are bi-allelic, since they can only be detected when they are ongoing and their signature vanishes from fully differentiated plasma cells or from “resting” blood memory B-cells, but readily reappears when such memory B-cells are re-stimulated *in vitro*. Highly diversified breakpoints are distributed either within the upstream (3’RR1) or downstream (3’RR2) copies of the IgH 3’ super-enhancer and all conditions activating CSR *in vitro* also seem to trigger LSR.

**Author Summary:** Class switch recombination, initiated by the activation-induced deaminase enzyme rearranges immunoglobulin (Ig) genes in order to replace expression of IgM by IgG, IgA or IgE. A variant form of this event, locus suicide recombination (LSR), was previously reported in mouse B-lymphocytes and simply deletes all functional Ig constant genes, thus terminating B-cell function. This study first demonstrates that the structure of the human Ig heavy chain locus provides an ideal target for LSR, and is thus actively (but transiently) affected by this deletional process at the activated B-cell stage. LSR then yields recombined genes that do not support B-cell survival and which thus become undetectable among long-lived memory B-cells or plasma cells.

## Introduction

Humoral immune responses and immunoglobulin (Ig) production rely on the selection of B-cells harboring antigen (Ag)-specific B-cell receptors (BCRs). This selection implies not only proliferation and differentiation of those cells optimally binding Ag but also elimination of the less efficient or inappropriately activated cells. The latter can be accomplished through various pathways leading to anergy, death-by-neglect or activation-induced cell death (AICD). While AICD pathways have been characterized in detail for T-cells, and notably involve FAS-induced apoptosis, they are less documented in B-cells. A major and unique feature of mature B-cells during Ag-driven responses, is their ability to reshape their genome, and more specifically Ig genes, after activation-induced deaminase (AID)-dependent modifications. Somatic hypermutation (SHM) within germinal centers (GC) can yield cells with higher affinity V domains, which preferentially capture Ag from follicular dendritic cells, undergo stimulatory cognate interactions with T follicular helper cells and are further selected for survival. In parallel, AID-dependent class switch recombination (CSR) diversifies IgH classes by joining repetitive switch (S) regions that precede the various constant (C) genes. Besides the selected winners of AID-mediated reshaping, many cells are losers or undesired responders deserving elimination, in agreement with the considerable amount of GC B-cells which have been demonstrated as actively undergoing apoptosis [1,2]. Out-of-frame or other unfavorable V region mutations might result in BCR loss and promote apoptosis while more subtle cell fate decisions will arbitrate between death, short-term or long-term survival as memory lymphocytes or plasma cells [1,3]. Such intra-GC cell fate choices are crucial since inappropriate survival or terminal differentiation of bystander cells producing useless Ig or Ig with increased affinity for self or environmental Ags might trigger auto-immunity, inflammation and disease. Since class-switched antibodies are potent actors of auto-immunity and/or hypersensitivity, means for restricting CSR and reentry of class-switched cells into SHM thus appear as necessary safeguards to keep humoral immune responses both specific and innocuous. The AID-dependent process of locus suicide recombination (LSR) reported in mouse B-cells ideally fits this necessity [4]. We now show that the human IgH 3’RR super-enhancers include sequences ideally suited as recombination targets, and that LSR actively occurs in human lymphoid B-cells.

LSR features recombination between Sμ and the 3’ regulatory region (3’RR) located downstream of the IgH locus, thus deleting the whole IgH constant gene cluster. The 3’RR includes several B-lineage specific enhancers (hs3, hs1–2 and hs4) which are mostly active after B-cell activation and then promote transcription and recombination [5]. Targeted mutations of the 3’RR in the mouse demonstrated its major role in SHM, its control of germline transcription and CSR to most C genes, its booster effect on IgH gene expression in plasma cells and even its ability to promote inter-allelic transvection between IgH alleles [5–11]. The mouse 3’RR has a unique palindromic architecture with functional implications [12–15], and this structure is shared by all mammalian species studied to date, including humans [16]. In addition, the mouse 3’ RR includes multiple stretches of repetitive DNA resembling S-regions albeit shorter [4,12], potentially promoting LSR through a mechanistic process similar to CSR. In B-cells, Ag stimulation with appropriate co-stimuli induces AID expression and then initiates staggered DNA nicking in repetitive S-regions, double strand breaks (DSBs) and CSR. Highly repetitive “Like switch” (LS) regions from the 3’RR might serve a similar role, ending with Sμ-3’RR junctions and deletion of the entire IgH constant region gene cluster. By abrogating BCR expression, which is critical for B-cell survival, this eliminates the corresponding cells.

In humans and old-world apes, an internal duplication of the IgH locus constant gene cluster has additionally duplicated 3’ regulatory regions downstream of each Cα gene (3’RR1 and 3’RR2respectively downstream of Cα1 and Cα2)[16,17]. We show that both human 3’RRs include LS-regions which are highly structured in terms of DNA repeats and which can locally be G-rich, with predicted ability to form the G-quadruplex (G4) DNA now known as the ideal substrate for AID [18,19]. Importantly, these regions also actively undergo AID-dependent recombination in activated B-cells, whatever the activation conditions assayed, both *in vivo* and *in vitro*.

## RESULTS

### Human 3’RR1 and 3’RR2 share multiple common features with S regions

Architectures of human 3’RR1 and 3’RR2 elements were analyzed for the presence of direct and inverted repeats (Fig1). As previously described for the mouse 3’RR, “like-switch” (LS) sequences (*i.e*. stretches of highly repetitive DNA somehow resembling S regions) were readily evidenced using the YASS algorithm (http://bioinfo.lifl.fr/yass/yass.php) for a survey of direct repeats in window range set over 20bp, with a minimal identity threshold set at 90% (Fig. 1A). Five to six human LS regions stand in each of the human IgH 3’RR super-enhancers. These LS regions are each 0.5 to 2 kb long, and are interspersed with core enhancer elements, which they do not overlap.

**Figure 1:**
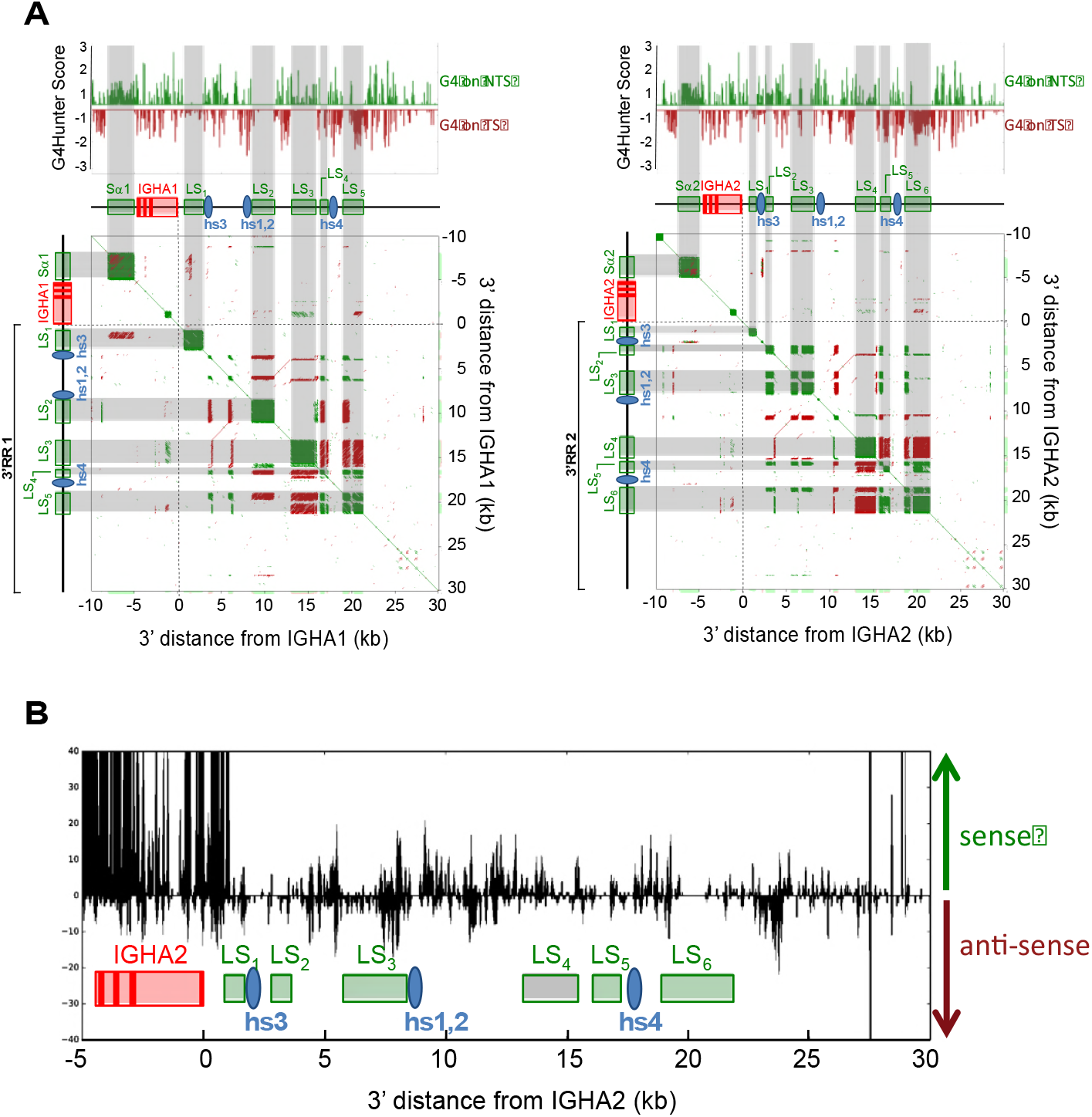
3’RR structure and transcription in activated B-cells. **A.** Structure of genomic fragments including either human Sα1, Cα1 gene and 3’RR1 (left), or Sα2, Cα2 and 3’RR2 (right) was analyzed for presence of G4 DNA on the template (TS) or non-template (NTS) strand (***Top***). Dot plot analysis of internal repeats (***Bottom***) was ran with the YASS algorithm: colored dots indicate short stretches of homology in direct (green) or inverted (red) orientation (green squares thus mark positions of highly repetitive S or LS regions / red lines indicate positions of inverted repeats, notably the palindrome centered on the hs1,2 enhancer). **B**. Transcription of human 3’RR2 in sense and antisense orientation, as evaluated by RNA-seq experiments with RNA from *in vitro* activated B-cells.

Human 3’RR LS sequences additionally show 35–50% G-richness on one or the other DNA strand, and analysis using the G4-hunter algorithm http://bioinformatics.cruk.cam.ac.uk/G4Hunter/ [23], predicted the occurrence of several clusters of G4-DNA, standing in either one or the other DNA strand of LS regions (Fig 1B).

When checking RNA-seq data from activated human B-cells, we noticed that 3’RR transcription occurred in both the sense and antisense orientation in reference to IgH constant genes, and that, quantitatively, it revealed equivalent amounts of transcripts corresponding to either orientation (Fig. 1B). This is in contrast with IgH constant genes (see for example Cα and the preceding Sα region on Fig 1B), which are predominantly transcribed in the sense orientation.

### LSR occurs at multiple sites within the whole extent of both human 3’RR

As previously described for mouse LSR, we identified human LSR junctions by nested PCR in DNA from B-cell samples [4]. The specificity of the read-out for these PCR assays was improved by replacing the final hybridization step of PCR products, initially done with specific probes[4]), with high-throughput sequencing adapted from our recently described CSR evaluation algorithm, CSReport [22]. Sequence determination thus precisely mapped the breakpoint sites. CSReport mapped LSR junctions linking Sμ to either the human 3’RR1 downstream of Cα1 or 3’RR2 downstream of Cα2.

Although such an “LSR-seq” DNA amplification method (with some amplified fragments above 30kb) is necessarily biased towards amplification of the shorter fragments, the use of 3 primers located either downstream from the h3, hs1,2 and hs4 enhancers finally amplified breakpoints located throughout 3’RR1 and 3’RR2.

These breaks were joined to various positions within Sμ, with preferential use of breakpoints at the 5’ side of Sμ, especially for junctions with the downstream part of the 3’RRs. When considering the sum of all junctions mapped in this study (Fig. 2), the most striking observation was their highly diversity and scattered distribution throughout the 3’RRs (both within LS regions and in their proximity).

**Figure 2:**
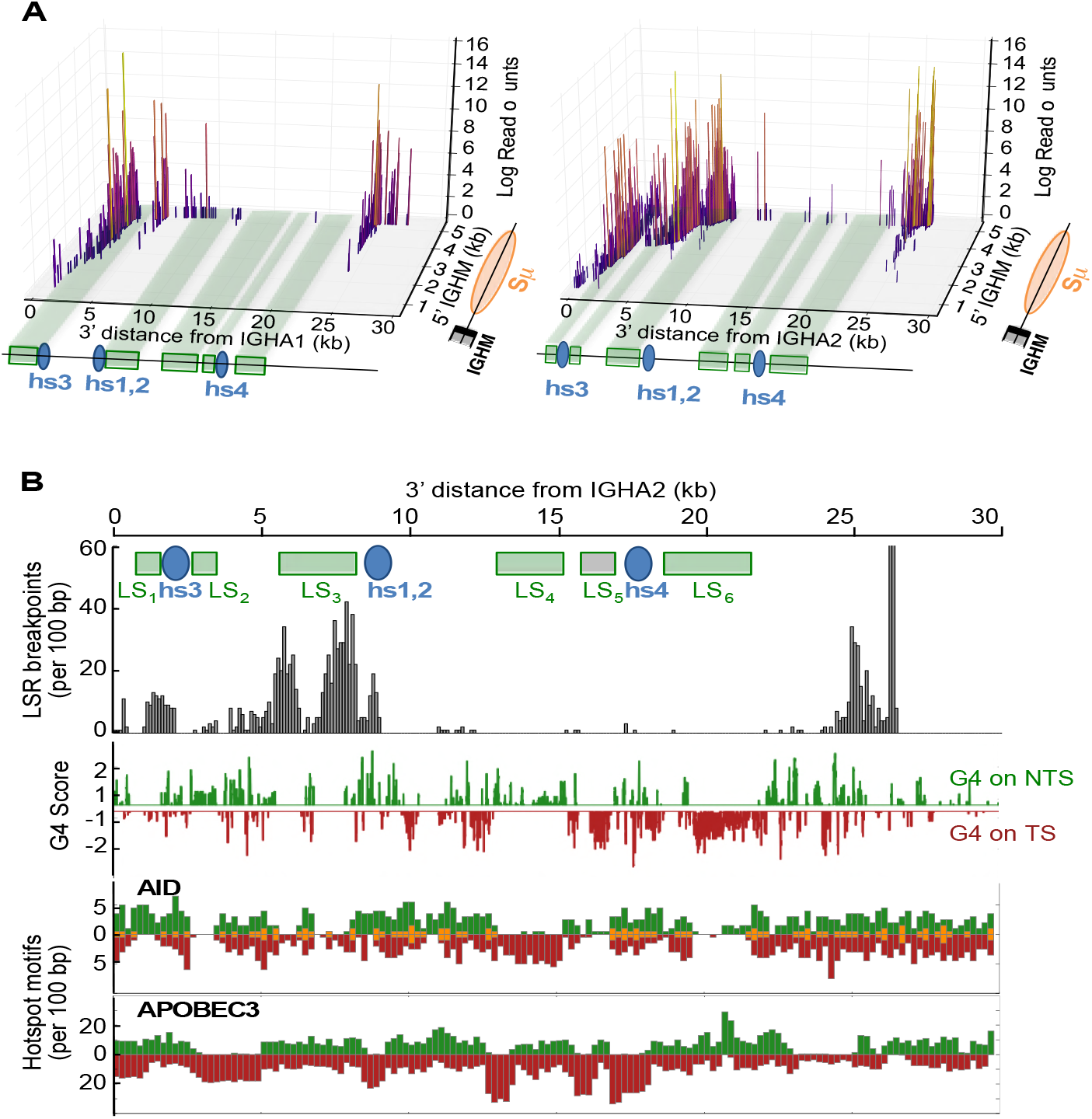
Positions of LSR breaks in human activated B-cells. A. **A 3D**-schematic showing the positions of junctions sequenced in this study with regards to abundance among sequence reads (in log scale) and position of DSBs in Sμ, and 3’RR1 (left) or 3’RR2 (right). **B**. Map of 3’RR2 reporting breaks observed in sequenced LSR junctions together with the position of G4 DNA on either the non-template (green) or template (red) strand (with reference to the transcriptional orientation of IgH constant genes), AID consensus target sites, and APOBEC3 family deaminases.

It is questionable whether LSR targets only unswitched B-cells, determining the occurrence of Sμ-3’RR junctions, or can also target class-switched cells and then eventually conserve a remnant of a downstream S region inserted in-between Sμ and a recombined 3’RR. We indeed obtained multiple evidence of such “post-switch” LSR, with sequence reads obtained from the LSR-seq protocol (*i.e*. sequencing an Sμ to 3’RR amplicon) but including junctions of Sμ with Sγ or Sα, (or Sγ or Sα joined with a 3’RR). Within a total of 11179 LSR junctions sequenced in this study, 1126, *i.e*. about 10% could be typed as complex junctions involving breaks with a downstream S (Sγ or Sα) region and certifying their origin from switched cells. Conversely, direct Sμ-3’RR junctions could originate from either unswitched or switched cells (upon complete deletion of the intervening S regions), so that the sequence of a given LSR junction cannot by itself certify that it occurred in an IgM+ cell.

### LSR-seq can semi-quantitatively estimate *in vivo* LSR in freshly collected B-cells

CSR junctions were characterized in parallel to LSR junctions, by a similar PCR/sequencing strategy in order to provide an LSR/CSR comparison. “LSR-seq” and “CSR-seq” protocols thus provided two semi-quantitative assessments of recombination events: numbers of <u>sequencing reads</u> including the expected junctions, and numbers of <u>unique junctions</u>. Although we cannot claim such a nested PCR and sequencing assay to be fully quantitative, it appeared to be at least semi-quantitative, giving negative results in non-B-cell samples and abundant signals in B-cells from chronically inflamed lymphoid tissues. It also had the abovementioned advantage of formally identifying sequencing reads mapped as CSR or LSR junctions, and allowed us to count repeated reads with identical sequences as “unique junctions”.

LSR junctions from Sμ to either 3’RR were accordingly “semi-quantified” in DNA from fresh primary B-cells from various origins. Peripheral blood mononuclear cells (PBMC) from healthy donors regularly yielded low, close to negative, LSR signals, and the same was observed in a series of patients recently treated for bacterial infections. By contrast, LSR was abundant in DNA from tonsils or adenoids. No LSR junctions were found in DNA from either tonsil B-cells nor PBMC obtained from AID-deficient patients. In healthy donors, as expected from the weak signals from PBMC, LSR was also absent or very low in sorted [CD19+, CD27+] peripheral blood memory B-cells and in sorted bone marrow CD138+ plasma cells (Fig. 3). LSR junctions were thus rare in DNA from circulating B-cells of healthy subjects and in differentiated plasma cells or memory cells. Further splitting and sorting of tonsil cells showed that LSR junctions occurred at a high level in cells with ongoing activation (sorted as blasting CD19+IgD-CD10-CD38^hi^ B-cells) (Fig. 3).

**Figure 3:**
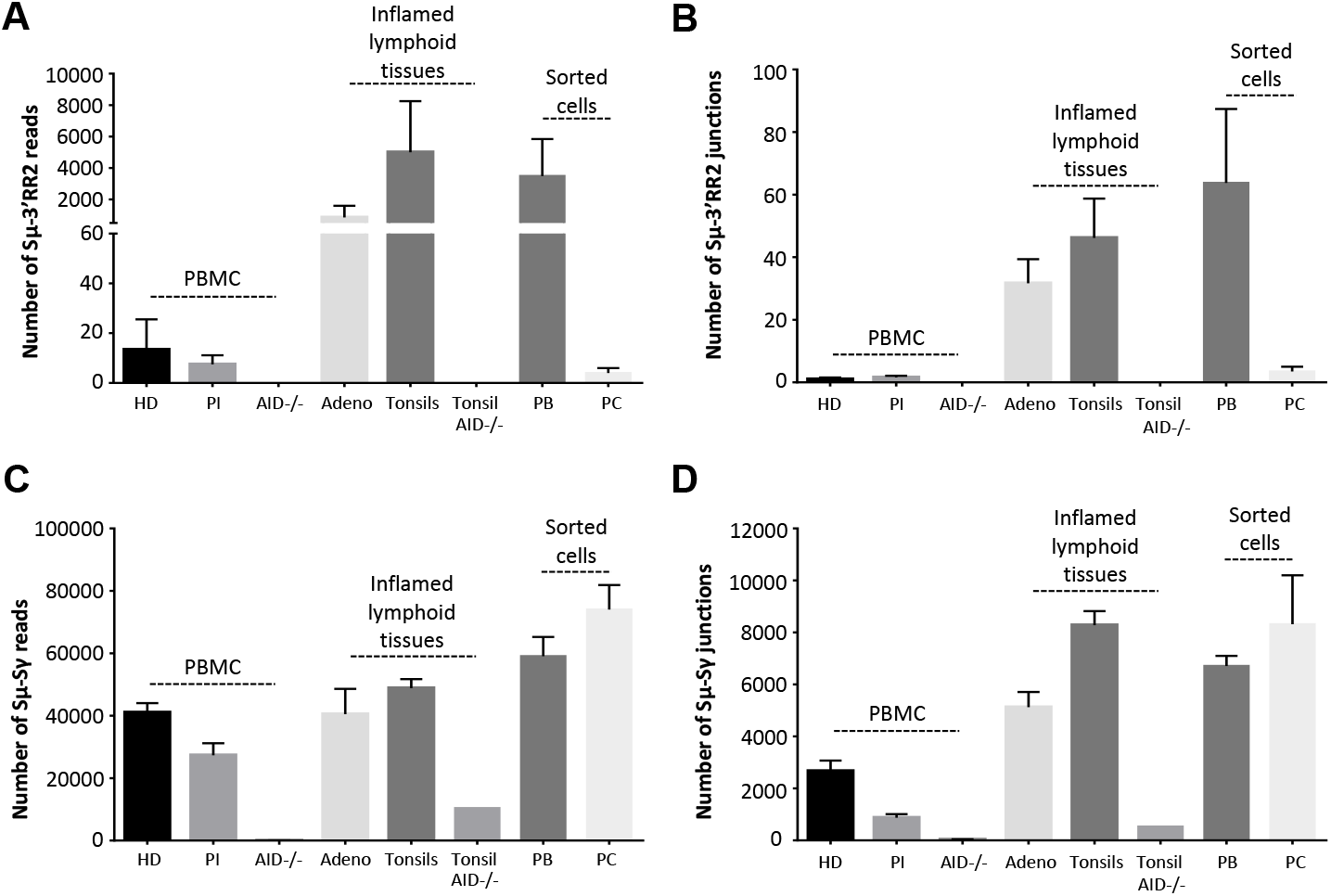
LSR breaks *in vivo* in humans. Sμ-3’RR2 LSR reads (**A**) and junctions (**B**) were analyzed in peripheral blood B cells isolated from healthy donors (HD) (n=6) but also in blood from post-infection (PI) patients recently hospitalized and treated for bacterial infections (n=17) and from AID-/- B-cells (n=3). LSR was also studied using DNA from inflamed lymphoid tissues (adenoids (n=3) and tonsils (n=11)), 1 AID-/- tonsil, from sorted plasmablasts (PB) isolated from tonsils (n=5) and finally in DNA from terminally differentiated plasma cells (PC) (n=2) sorted from bone marrow. Frequencies of LSR junctions amplified with an Sμ forward primer and a downstream Hs4 reverse primer were evaluated in parallel to those of classical Sμ-Sγ CSR reads (**C**) and junctions (**D**) amplified from the same DNA samples, in order to provide an LSR/CSR comparison, data represent mean numbers (reads or junctions) ± SEM.

### LSR junctions actively occur *in vitro* in activated B-cells

Class-switching and induction of AID in human B-cells is mostly a T-dependent process and is most often modelled *in vitro* by activating B-cells with CD40 ligands. When total blood B-cells from healthy patients were stimulated in such conditions, this resulted in the induction of LSR simultaneously with CSR. Associating anti-CD40 stimulation with various co-stimulatory factors, increased LSR and the LSR/CSR ratio in most of the conditions assayed and notably upon addition of IL4, IL21, or BCR ligation with anti-κ antibodies, or, to the highest extent, upon TLR ligation (Fig. 4).

**Figure 4:**
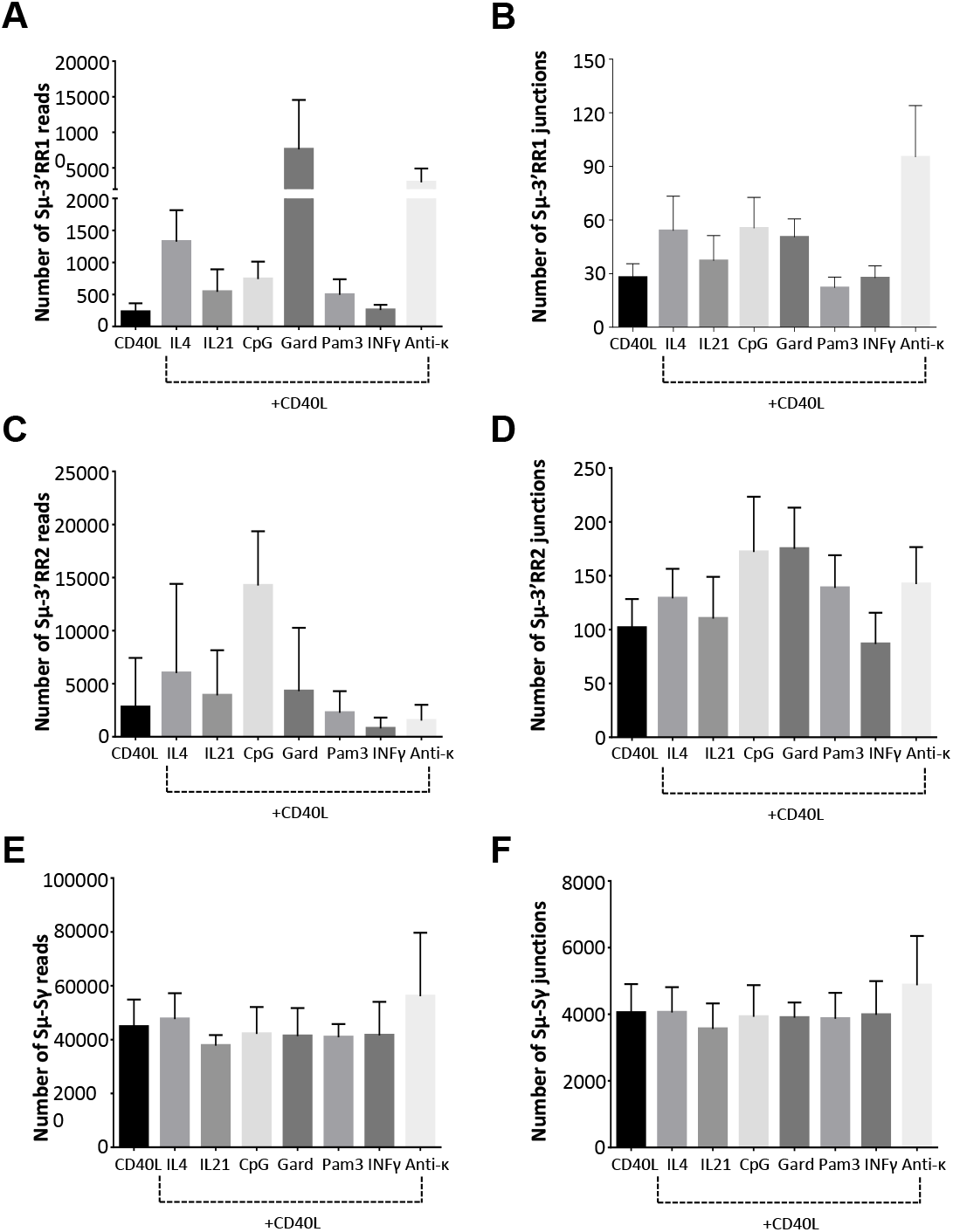
LSR after *in-vitro* stimulation of peripheral blood B-cells in various conditions of activation. PBMC from healthy donors activated for 4 days with human recombinant CD40L alone or with recombinant human IL4, IL21, CpG, gardiqimod (Gard), Pam3, IFNγ or with an anti-Igκ antiserum. Sμ-3RR1 (**A** and **B**) and Sμ-3RR2 (**C** and **D**) LSR junctions were amplified using reverse primers located in hs3, hs1–2 and downstream hs4. Equal amounts of the three PCR products were mixed in a single library. **E** and **F** represent IgG CSR reads and junctions using the same stimulation conditions. Data represent means (for unique reads and junctions) ± SEM for 6 people.

We also sorted three different activated B-cell stages, using a 6-day long *in vitro* activation protocol with combined anti-CD40, TLR9 ligand CpG, anti-BCR antibodies and cytokines, previously described as optimal for mimicking the *in vivo* humoral immune response [21] (ref). This allowed us to generate either plasmablasts committed towards plasma cell differentiation, or activated B-cells having proliferated but still at an earlier stage (resembling proliferating centroblasts), and to compare them to surviving but non-dividing B-cells (bystanders) from the same culture. Analysis of the three types of cells showed that LSR was virtually absent in non-dividing bystander B-cells, but was, by contrast, at the highest level in plasmablasts and at an intermediate level in actively proliferating activated B-cells (Fig. 5A).

**Figure 5:**
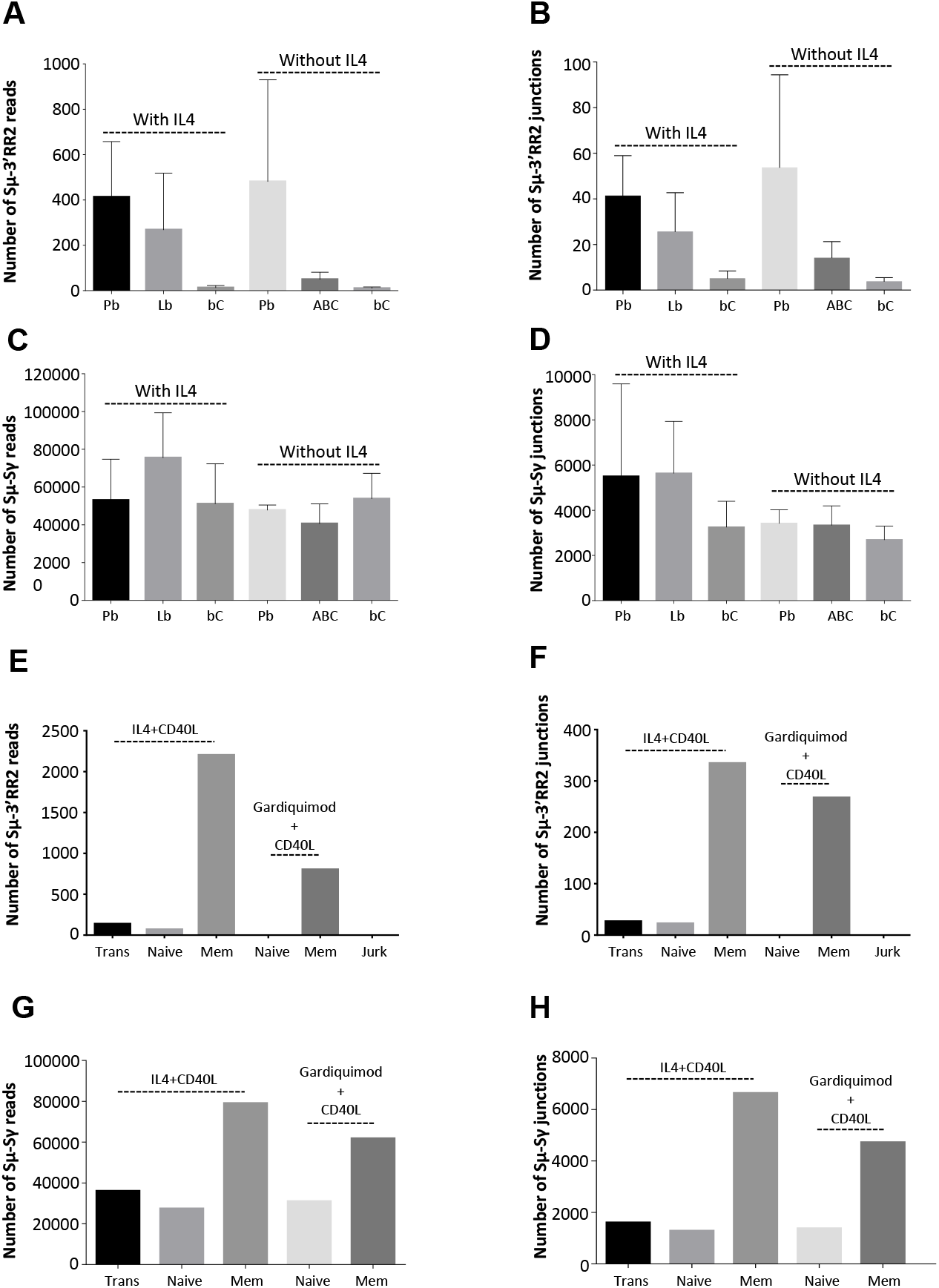
LSR breaks in vitro in stimulated B-cells from various origins. **A** and **B**. LSR (reads and junctions) in *in vitro* stimulated naive B-cells, sorted after stimulation. PBMC from healthy donors were activated *in-vitro* for 6 days with combined anti-CD40, TLR9 ligand CpG, anti-BCR antibodies and other cytokines generating three different stages of differentially activated-B-cells: Pb (plasmablasts), ABC (proliferating activated B-cells) and bC (bystander B-cells). Data represent mean number (reads or junctions) ± SEM for 3 donors. **C** and **D**. CSR (reads and junctions) to Sγ in same samples as A and B. CSR junctions were amplified using a downstream IgG consensus primer located in CH1. **E** and **F**. Reads and junctions from *in vitro*-induced LSR in stimulated B-cells from various origins: sorted transitional (Trans), naive and memory B-cells (Mem) (n=1) B-cells obtained from peripheral blood of healthy donors. Before DNA extraction, cells were stimulated for four days *in-vitro* with human recombinant CD40L combined with human recombinant IL4 or with gardiquimod. Jurkat (Jurk) cells stood as negative controls. **G** and **H**. CSR reads and junctions from same samples as in E and F. CSR junctions were amplified using a downstream Cγ consensus primer located in CH1. Data represent means (for unique reads and junctions) ± SEM.

We also studied LSR in cells first sorted and then stimulated *in vitro*, comparing memory, naive and transitional B-cells (sorted as CD27+CD38^lo^CD24^lo^, CD27-CD38^lo^CD24^lo^ and CD27-CD38^h^iCD24^hi^ subsets, respectively). We found that upon *in vitro* activation, memory B-cells yielded much higher levels of LSR than naive or transitional B-cells (LSR remaining undetectable in control Jurkat cells) (Fig. 5, E and F). As expected in sorted populations including from the beginning a large amount of switched cells, CSR junctions assayed in parallel were the most abundant in memory cells (Fig. 5, G and H).

### Structure of LSR junctions

CSR junctions from Sμ to Sγ and LSR junctions from Sμ to 3’RR were amplified by PCR with specific primers, sequenced by NGS (Ion Proton) and analyzed regarding their positions in S regions and/ or 3’RR, and their nature using CSReport, a dedicated tool that performs junction assembly and analysis based on BLAST alignments (Fig. 6). A first difference between LSR and CSR concerned the global position with regard to repetitive regions. All Sμ breaks were located in the repetitive part of Sμ (in agreement with the fact that in the mouse, deletion of Sμ repeats virtually abrogates CSR)[24]. By contrast in the 3’RRs, breaks were found not only within but also outside of LS regions. A second difference concerned the detailed position of 3’RR breaks (as identified in sequenced unique junctions) and their increased distance relative to the various genomic features relevant to DNA cleavage (main AID target sites WRCY and RGYW, G4 sites suggested by the G4 Hunter algorithm) (Fig. 6, A and B). Interestingly in LSR junctions, the increased mean distance concerned only 3’RR breaks, while Sμ breaks were similar to CSR junctions. These increased mean distances between breakpoints in unique junctions and relevant genomic features (in color) were compared to the same distance calculated from simulated random break positions in the sequenced regions. Even if the mean [break to AID site] distance was longer in the 3’RR than in Sμ, it remained significantly shorter than that calculated under the random break hypothesis. The same evaluation was done for APOBEC3 family target sequences, which by contrast did not show any difference between the observed distance and that calculated under the hypothesis of random breaks (Fig. 6C).

**Figure 6:**
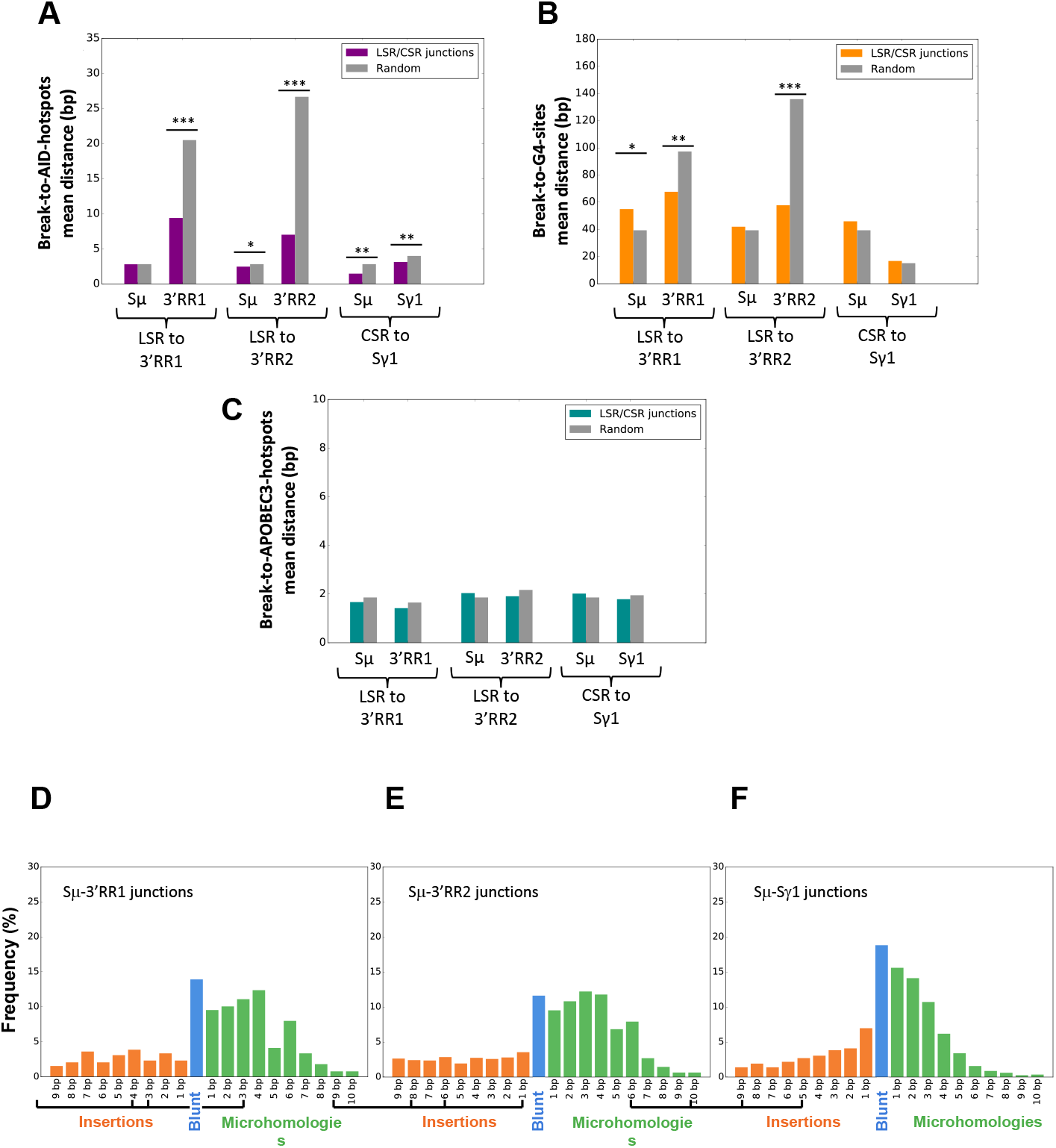
Detailed position of breaks and structure of human repaired LSR junctions. Mean distance to AID target sites (**A**), G-quadruplex (**B**) and APOBEC3 family (**C**) sites. Structure of repaired junctions with regards to mean number of inserted nucleotides and length of microhomology between broken ends for Sμ-3’RR1 (**D**), Sμ-3’RR2 (**E**) and Sμ-Sγ1 (**F**).

Beyond DNA breaks, another difference between CSR and LSR was related to the junctional sequence itself. Our CSReport algorithm indeed classifies junctions as either blunt (broken ends directly joined without overlap), with microhomologies (stretches of a few nucleotides shared between joined segments) or with insertions (a few nucleotides inserted between joined segments). Figures 6, D-F compare more than 2000 LSR unique direct junctions characterized in this study (≈ 20% Sμ-3’RR1 and ≈ 80% Sμ-3’RR2 junctions) to a thousand of independent unique Sμ-Sγ1 junctions obtained from one tonsil sample. LSR and CSR somehow differed: LSR to both Sμ-3’RR1 and Sμ-3’RR2 exhibited a lower frequency of blunt junctions (12 % *vs*. 19%) and longer microhomologies (mean length 3.8bp *vs*. 2.7bp) and insertions (mean length 4.8bp *vs*. 3.7bp).

## DISCUSSION

Analyzing the architecture of human IgH 3’RR elements shows their high architectural similarity with the mouse 3’RR, notably for the presence of a central palindrome centered on the hs1,2 enhancer and for the presence of several stretches of repetitive LS regions. The latter LS regions are each 0.5 to 2 kb long, which is close to the length of Sα2 and appears longer than LS regions in the mouse 3’RR, with the longest LS flanking the hs4 core enhancer. Although in lower amounts than for S-regions, human 3’RR1 and 3’RR2 also include G-rich DNA, but not restricted to a single DNA strand. A specific feature of mammalian S regions is indeed G-richness of the non-template strand, where it promotes the initiation or extension of R-loops, and also often determines tandem G-repeats with potential to form G4 DNA [18,19,25]. In S regions, G4-DNA promotes the occurrence of DNA nicks and further excision by exonuclease 1, while pharmacological agents binding G4-DNA can inhibit CSR [18,26,27]. I*n vitro*, G4-DNA was also shown to provide a type of structured DNA which favors the activity of Activation-Induced Deaminase (AID) [19]. Finally, G-quadruplexes forming in S region transcripts were shown to recruit the helicase Ddx1 and contribute to the formation of R-loops before CSR [28].

The G4-hunter algorithm also identified G4-DNA forming sequences within the human 3’RRs, together with long G-rich stretches standing in either one or the other DNA strand. In this regard, it should be noticed first that human 3’RR transcription occurs in both the sense and antisense orientation (with reference to *IgH* constant genes) and secondly that 3’RR eRNA are non-coding transcripts, for which definition of a “template strand” (as for *IgH* constant genes) is ambiguous. In such settings, G4 structures from any DNA strand might recruit AID activity within the 3’RR LS regions, similarly to G4-DNA from the non-template strand within S-regions.

The present study demonstrates a high diversity of IgH recombination events joining Sμ or downstream S regions to the 3’RR super-enhancers in activated human B-cells from healthy individuals, showing inter-species conservation of the LSR process previously observed in the mouse. This process appears to be AID-dependent since such junctions were virtually absent in tissues from AID-deficient patients. Study of cells from various origins indicated that human LSR junctions could only be detected transiently when cells were activated but were not preserved in the long term in quiescent memory B-cells. Indeed, when DNA from peripheral blood B-cells was collected from various individuals or from bone marrow plasma cells, it gave low or negative LSR signals by LSR-seq, while CSR signals were high. Since peripheral blood is known to include one-third of memory B-cells in adults [29], this observation suggests that IgH alleles carrying LSR deletions are absent (or exquisitely rare) in circulating memory B-cells and thus that they rarely target the non-functional allele. Rather, LSR deletion likely predominantly targets the functional IgH allele (or is biallelic), and therefore incompatible with further survival of B-cells in the absence of a functional BCR. On the contrary, a major portion of circulating memory B-cells obviously survive class-switching and accounts for persisting CSR junctions detectable in DNA from PBMC [29].

By contrast to blood, human lymphoid tissues, known as sites of chronic immune stimulation, such as tonsils and adenoids, yielded abundant LSR junctions by LSR-seq. Contrasting with spleen or blood, tonsils and adenoids are known to include abundant GC B-cells with an ongoing activation phenotype (CD20high, BCR low, often showing CD38 expression) [29,30]. B-cells from such chronically activated lymphoid tissues thus include a mix of naive cells, memory cells (about 20%) and GC activated B-cells as well as plasmablasts involved in ongoing local immune reactions to mucosal antigens.

This major difference between blood and tonsils thus confirms that LSR signals in humans are not associated with the presence of memory B-cells (by contrast to CSR) and indicates that LSR is only transiently detectable contemporaneously to *in vivo* B-cell activation.

Human LSR junctions were related to either 3’RR1 or 3’RR2 and disseminated throughout these regions. The main conclusion to be drawn concerning this distribution is the high diversity of the breakpoints. The apparent higher frequency of breaks at the beginning of Sμ likely relates to the preferential amplification of short fragments in a nested PCR assay. Breaks within the large 3’RR, also appeared to cluster in three sub-regions each spanning 3 to 5 kilobases, but these sub-regions in fact preceded the three unique non-repetitive locations where it had been possible to design highly specific reverse primers for the LSR-seq assay. While the use of additional regularly spaced reverse primers is prevented by the highly repetitive 3’RR sequence, it is thus likely that the frequency of breaks occurring at large distances from our primers is under-estimated.

Interestingly, sequence analysis of LSR-seq data from both *in vivo* and *in vitro* activated B-cells revealed the presence of both direct junctions from Sμ to one or the other 3’RR, or of complex junctions including Sγ or Sα regions in addition to Sμ.

Evaluation of LSR and comparison to CSR is inherently hindered by the need to identify junctions at the DNA level using nested PCR and sequencing (which can only be semi-quantitative), and by the fact that suiciding cells rapidly vanish while switched cells are healthy and CSR remains detectable in the long term. In spite of that, we routinely identified LSR reads in DNA samples from activated B-cells, with frequencies up to 3% of the detection rate for CSR reads, meaning the LSR is a frequent event.

LSR junctions were found both in cells activated *in vitro* and in cells activated *in vivo* after CD40 ligation. It was not possible to identify conditions stimulating CSR in the absence of LSR. Even if variations appeared, with a tendency to increase the LSR/CSR ratio when TLR ligands were associated with CD40 ligation, or to decrease this ratio when BCR cross-linking was associated, these variations remained below statistical significance.

Another interesting point is the mechanisms of DNA breaks occurring in the 3’RR. While the process of LSR is globally AID-dependent, we cannot exclude that breaks in Sμ might be initiated by AID while breaks in the 3’RR might be generated by another mechanism, yet to be identified. While the 3’RR includes repeats and is transcribed, similarly to a S region, our analysis showed that the distance between AID consensus motifs and 3’RR DSBs was much higher than for S regions. The same is true for the distance between breaks and G quadruplexes. This increase might in part relate to the structure of the target sequences: S is indeed extremely rich in AID hotspot motifs and G4 DNA; the mean distance of breaks to these target sites was accordingly very low (and did not differ significantly from the distance calculated under the hypothesis of random breaks). In contrast, both 3’RRs exhibit a lower density of AID hotspots and G4 DNA. 3’RR DSBs identified in unique junctions were located significantly closer to these sites than they would be if randomly distributed. This suggests active targeting of these sites by AID, even if the mean distance from AID sites is higher than for Sμ breaks (or for Sγ breaks in CSR junctions). While the similarity of Sμ breaks in CSR and LSR junctions strongly suggests that identical mechanisms are at work, the differences seen for the 3’RR might suggest that, in addition to the structure of the targeted sequence, either AID or its partners are differentially involved in both regions. The occurrence of breaks both within and outside LS repeats together with the lower density of AID sites suggest that, compared to Sμ, DSBs less easily follow staggered SSBs in the 3’RR and require more complex processing of initial SSBs. DNA cleavage outside of a repetitive target sequence has been previously reported for CSR to the Cδ gene, where the target Σδ sequence harbors few AID hotspots [31]. Even in classical S regions, once initiated in a repetitive region, R-loops eventually extend at distance though nearby G-rich regions, and hereby increases the length of the region targeted by CSR [25].

LSR was nearly absent in samples from AID-deficient individuals but we cannot exclude that this would only result from the absence of DSBs in Sμ. There is however no evidence indicating that enzymes other than AID might contribute to LSR breaks, and notably the evaluation of distance from breaks to APOBEC3 family targets sites more or less ruled out that these deaminases might significantly contribute to LSR. Altogether, the significant implication of AID hotspots despite their relative scarcity implies that, in the 3’RR, cytosine deamination is followed by DNA remodeling on longer distances than within Sμ before occurrence of DSBs, synapsis to an S region and repair. The hypothesis of a change in the structure of the broken ends fits quite well with the observed increased length of microhomologies in repaired junctions. LSR is indeed repaired less frequently than CSR with blunt junctions typical of non-homologous end-joining, but with an increased usage of microhomologies and longer insertions, as described in the situation with increased involvement of alternate end-joining and microhomology mediated repair.

AID-initiated locus suicide recombination of the IgH locus thus actively occurs in human B-cells, with downstream breakpoints throughout the 3’RR super-enhancer. (Fig. 7). The phenomenon either predominates on the functional IgH allele or is bi-allelic, since LSR deletions are only detected transiently during B-cell activation but do not remain detectable in memory cells or plasma cells.

**Figure 7:**
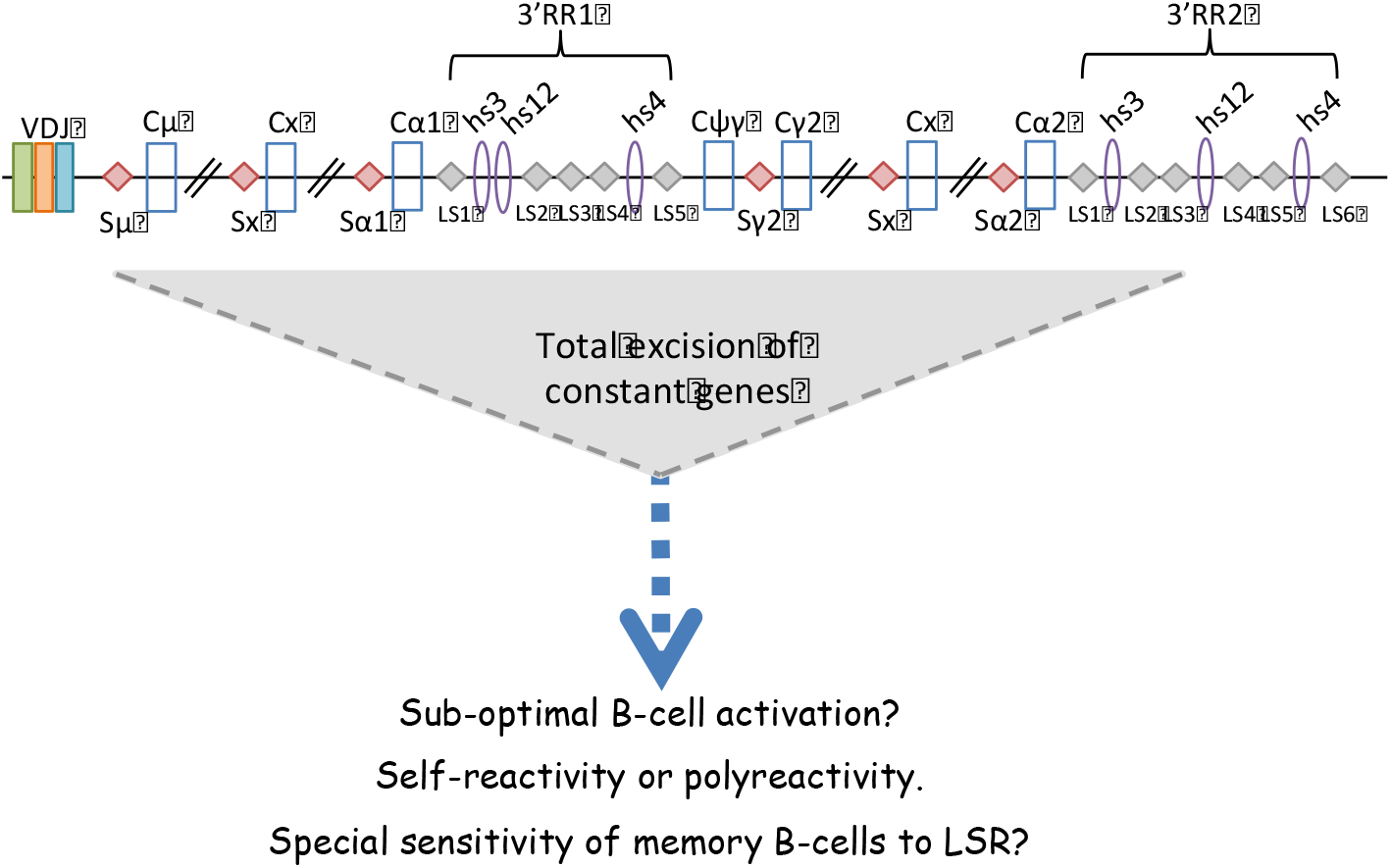
Model of LSR in the human IgH locus. Recombination between Sμ and LS regions in the 3’RR2 super-enhancer leads to complete deletion of constant regions of all immunoglobulin classes leading to absence of BCR membrane expression. LSR could be a physiological mechanism to eliminate sub-optimally activated B-cells or those that are poly or autoreactive. Resting memory B-cells seem especially sensitive as they do not carry stigmas of LSR, which only occurs in activated B-cells.

Because it is transient and followed by cell death, it remains difficult to evaluate LSR regulation and impact on B-cell homeostasis, relative to the other cell death pathways active in the GC. Few methods are available for studying deletions of hundreds kilobases with loosely determined breakpoints and the use of PCR followed with next generation sequencing remains semi-quantitative, given the PCR obstacles introduced by the repetitive nature of the 3’RR. In such settings, all conditions assayed for B-cell activation induce LSR at some degree and new methods will be necessary in order to determine where and when LSR occurs *in vivo* at the single-cell level.

## Acknowledgements

We are grateful to M. Brousse, M. Faïsse, S. Desforges and B. Remerand for technical help, and to Dr. Sven Kracker for help in the transfer of human samples.

## Authorship Contributions

MC designed the experiments, ID and ZD performed the experiments, AP, GL and TF provided B-cells from human lymphoid tissues and performed some in vitro activation assays. CD and FC provided RNAseq data from human activated B-cell samples. AD provided blood and lymphoid tissue samples from AID deficient individuals. FB performed *in silico* analysis of data and characterization of CSR/LSR junctions. JCM, SP, JCA and MC analyzed the data. JCM and MC wrote the manuscript.

## MATERIALS AND METHODS

### Human samples

Blood from healthy volunteers was collected, after obtaining written informed consent, from the CHU Dupuytren Hospital, Limoges, France (ANSM: 2012-A00630–430) or from the Etablissement Français du Sang (Rennes, France; French Ministry of Higher Education and Research approval: AC-2014–2315) according to the Declaration of Helsinki. Blood from septic patients was collected from the Hematology department, CHU Dupuytren. The study was accepted by the Ethics Committee of CHU Dupuytren (n° 119–2013-19). The trial was registered under ClinicalTrials.gov: NCT01995448. Tonsils were obtained from patients undergoing tonsillectomy performed in CHU de Rennes, while adenoids were obtained from CHU Dupuytren. All were operative pieces. Bone marrow aspirates obtained from patients undergoing cardiac surgery. Subjects were recruited under institutional review board approval and informed consent according to the Declaration of Helsinki. Samples from AID-/- patients (3 PBMC samples and 1 sample of sorted tonsil B-cells) were provided by Pr Anne Durandy, Hôpital Necker, Paris, France.

### Activation of human B-lymphocytes *in vitro*

Human PBMC were obtained by density gradient centrifugation and human B-cells were negatively sorted from PBMC using EasySep Human B-cell Isolation Kit (StemCell) according to the manufacturer’s instructions. Purified B-lymphocytes were seeded at 1 × 10^6^ cells/ml in IMDM medium (Lonza) supplemented with 20% fetal bovine serum (Deutscher), 1% penicillin/streptomycin (Gibco) and stimulated for 4 days with 100 ng/ml human recombinant CD40L (<u>Enzo Life Sciences</u>) alone or with 50 ng/ml recombinant human IL-4 (Peprotech) or 2 μg/ml Gardiquimod (InvivoGen) or 2.5 μg/ml CpG oligodeoxynucleotide 2006 (InvivoGen) or 100 ng/ml IL-21 (R&D Systems) or 100ng/ml INFγ (R&D Systems) or 50 ng/ml Pam3CSK4 (InvivoGen) or 0.5 μg/ml goat anti-human kappa (Southern Biotech).

### Sorting of memory, transitional, and naive B-cells, and generation of *in vitro* activated plasmablasts from cultured naive B-cells

PBMCs from healthy volunteers were obtained after density centrifugation. Naive B-cells (NBCs) were purified by negative selection using magnetic cell separation (Naive B-cell Isolation Kit II; Miltenyi Biotec), following the manufacturer’s instructions, with the AutoMACS deplete-sensitive program. Purity of isolated CD19+CD27− NBCs was routinely >99%. Purified NBCs were labeled with 1 μM CFSE (Thermofisher) at 37°C for 10 min and washed in complete medium consisting of RPMI 1640 supplemented with 10% FCS and antibiotics (all from Gibco, Thermofisher). Purified human NBCs were cultured at 7.5 × 10^5^ cells/ml in 24-well plates and stimulated during 4 days with 2.6 μg/ml F(ab′)2 goat anti-human IgA+IgG+IgM (H+L) (Jackson ImmunoResearch Laboratories, West Grove, PA), 100 ng/ml recombinant human soluble CD40L (NCI), 2 μg/ml CpG oligodeoxynucleotide 2006 (Miltenyi Biotec), and 50 U/ml recombinant IL-2 (SARL Pharmaxie). Day-4 activated B-cells were washed and cultured at 4 × 10^5^ cells/ml with 50 U/ml IL2, 12.5 ng/ml IL-10 and 5 ng/ml IL-4 (R&D Systems). At day-6, cells were separated by cell sorting (FACSAria cell sorter, BD Biosciences) into undifferentiated CFSE^hi^ CD38^lo^ (bystander lymphocytes, without proliferation), CFSE^lo^ CD38^lo^ (proliferating activated B-cells), and differentiated CFSE^lo^ CD38^hi^ (plasmablasts) subsets[20].

After B-cell isolation (B-cell isolation kit II, Miltenyi Biotec), we also purified transitional, naive and memory B-cells which were sorted as CD27+CD38^lo^CD24^lo^, CD27-CD38^lo^CD24^lo^ and CD27-CD38^hi^CD24^hi^ subsets, respectively. Isolated cells were stimulated 4 days with different combinations of 100 ng/ml CD40L, 50 ng/ml IL4 and 2 μg/ml Gardiquimod (InvivoGen).

### Sorting of *in vivo* differentiated early plasmablasts

Actively differentiating plasmablasts from tonsils were purified as previously described [21]as CD19+IgD-CD10-CD38^hi^ cells. CD38^hi^/CD138+ mature plasma cells were purified from bone marrow aspirates.

### RNAseq on activated human B-cells

Human naive B-cells were purified and activated *in vitro* as described (Hipp et al. Nat Commun. 2017). RNA extractions were performed on day 4 IL-2 primed cells with the NucleoSpin RNA XS kit (Macherey-Nagel). Libraries were prepared with the TruSeq Stranded mRNA Library Prep Kit (Illumina) and samples were sequenced on an Illumina NextSeq 500 using 75-bp single-end reads (NextSeq 500 High Output v2, Illumina) by Helixio (Clermont Ferrand, France). Quality of sequencing data was monitored by FastQC. Residual adapters from sequencing were trimmed using Cutadapt 1.0. Potential PCR duplicates were removed using SAMtools 1.3. Reads were then aligned on the GRCh38 human genome using STAR 2.4.2a.

### Amplification of Sμ/Sγ for CSR-seq experiments

DNA from B-cells was extracted using <u>GenElute Mammalian Genomic DNA Miniprep K</u>it (Sigma). Sμ/Sγ junctions were amplified in triplicate by nested PCR with 200 ng DNA (<u>Herculase II Fusion DNA Polymerase, Agilent Genomics</u>) using the following consensus primers which amplify all four classes of human IgG junctions: PCR1: Sμ1a 5’- CCAGGTAGTGGAGGGTGGTA-3’ and IgGa consensus reverse 5’- GGTCACCACGCTGCTGAG-3’, PCR 2: Sμ1b forward 5’- CAGGGAACTGGGGTATCAAG-3’ and IgGb consensus reverse: 5’- CTTGACCAGGCAGCCCAG-3’. Forward primers used in both PCRs are located in Sμ while the reverse primers are located in the IgG consensus sequence of the CH1 region.

### Amplification of Sμ/3’RR junctions for LSR-seq experiments

Sμ/3’RR junctions were amplified under the same conditions as for Sμ/Sγ junctions using different primers located at different positions within the 3’ RR. The forward primer located in the Eμ region (EμH1) was used with a reverse primer located downstream of Hs4 (3’FarHhs4 reverse) or with a primer located in Hs1.2 (Hs1.2 reverse1) or in Hs3 (Hs3 reverse 1). PCR1 product was used to perform PCR2 using Sμ1b forward with all reverse primers located in 3’ RR. The primers sequences and PCR programs are given in Table I.

**Table 1.**
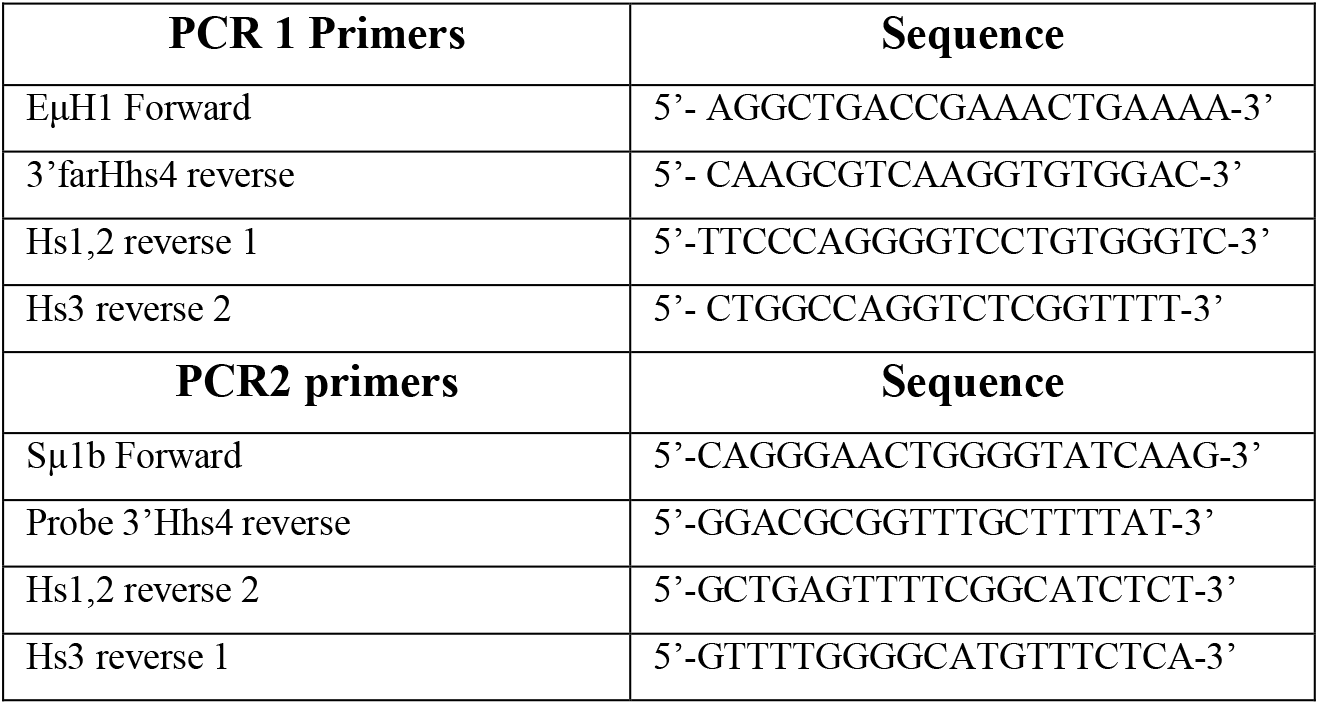
Primers used in nested PCR for amplification of LSR junctions

### Ion torrent next generation sequencing NGS

Barcoded libraries with 200-pb read lengths were prepared using Ion Xpress plus Fragment Library Kit (Thermo Fisher Scientific) according to manufacturer’s instructions. Each library was prepared using 200 ng PCR2 product. For LSR junction sequencing equal amounts of each PCR2 product (using hs3, hs12 and h4 primers) were mixed together and diluted to 100 pM to prepare one barcoded library. Libraries were run on an Ion PI v3 chip on the Ion Proton sequencer (Life Technologies). Data analysis was performed using CSReport [22].

### Data analysis

Data was analyzed by using GraphPad Prism software. Data are shown as means ± SEM of the indicated number of values. The student T test was used to determine significance between stimulation conditions.

